# The biophysical nature and not only the size of protein aggregates determines the strength of the immune response against dengue ED3

**DOI:** 10.1101/2022.11.02.514810

**Authors:** Md. Golam Kibria, Yukari Shiwaku, Subbaian Brindha, Yutaka Kuroda

## Abstract

Here we used domain 3 of dengue virus serotype 3 envelope protein (D3ED3), a natively folded globular low-immunogenicity protein, to ask whether the biophysical nature of amorphous aggregates can affect immunogenicity. We prepared amorphous oligomers in five distinct ways. One oligomer type was produced using our SCP tag (Solubility Controlling Peptide) made of 5 Isoleucines (C5I). The others were prepared by miss-shuffling the SS bonds (Ms), heating (Ht), stirring (St), and freeze-thaw (FT). Dynamic light scattering showed that all five formulations contained oligomers of approximately identical sizes with hydrodynamic radii (*Rh*) between 30 and 55 nm. Circular dichroism (cd) indicated that the secondary structure content of oligomers formed by stirring and freeze-thaw was essentially identical to that of the native monomeric D3ED3. The secondary structure content of the Ms showed moderate changes, whereas the C5I and heat-induced (Ht) oligomers exhibited a significant change. Immunization in JcL:ICR mice showed that both C5I and Ms significantly increased the anti-D3ED3 IgG titer. Ht, St, and FT were barely immunogenic, similar to the monomeric D3ED3. Cell surface CD marker analysis by flow cytometry confirmed that immunization with Ms generated a strong central and effector T-cell memory. This result adds a new dimension to earlier studies where the strength of the immune response was associated solely with the presence and sizes of the oligomers. It also suggests that controlled oligomerization can provide a new, adjuvant-free method for increasing a protein’s immunogenicity, yielding a potentially powerful platform for protein-based vaccines.

**Significance:** Protein aggregation is suspected to increase the immunogenicity of proteins. Here we show that the strength of the immune response depends not merely on the size of the oligomers/aggregates but also on their biophysical properties. Dengue virus 3 envelop protein domain 3 (D3ED3) was oligomerized/aggregated in five different ways. All five formulations contained oligomers with hydrodynamic radii between 30 and 55 nm. Two formulations, where D3 ED3 was natively folded, were not or poorly immunogenic. On the other hand, two others, where D3ED 3 was in a molten globule-like state, were strongly immunogenic. This result adds a new dimension to earlier studies where the strength of the immune response was associated solely with the presence and sizes of the oligomers.

## Introduction

Dengue virus (DENV) is a flavivirus with four distinct serotypes (DENV1-4). It is a positive-sense single-stranded RNA virus. The genome encodes three structural proteins (capsid (C), pre-membrane (prM), and envelope (E) protein) and seven non-structural proteins [1]. Domain 3 of the envelope protein (ED3) is attracting much attention owing to its involvement in host cell receptor interaction for host infectivity [2]. As such, it is the target of intensive research for developing antiviral drugs and vaccines [3].

Protein-based or subunit vaccines would have many advantages over the traditional whole-pathogen vaccine in production, handling and storage, and even safety [3–4]. However, unlike inactivated or live-attenuated whole pathogens, which possess intrinsic immune-stimulatory capable of inducing long-term protective immunity, proteins, especially small ones, are poorly immunogenic [5]. Subunit vaccines often use adjuvants to stimulate the immune response and enhance their efficacy as a vaccine antigen [6], but searching for new and safer adjuvants is time-consuming [7].

A rationale for the low immunogenicity of small proteins is their lack of pathogen-associated molecular patterns (PAMPs), a repetitive conformational pattern of the pathogen [8]. New approaches that use external substances to increase the antigen’s immunogenicity by producing repetitive conformational epitope patterns using biodegradable polymers [9] or nanoparticles [10] are being investigated. In particular, virus-like particle (VLP) is a recent strategy for enhancing a protein’s immunogenicity [11], which acts by presenting multiple copies of antigenic epitopes on its surface and thus mimicking a virus surface [12].

The amorphous aggregation of therapeutic proteins has long been suspected of inducing unexpected immune responses [8,13]. Besides several clinical observations where conditions are not well controlled, laboratory experiments show that oligomerization and aggregation of a protein might increase immunogenicity [14,15]. However, in most experiments, except for their sizes, the biophysical and biochemical nature of the oligomers/aggregates are not monitored. It thus remains unclear what kind of oligomers/aggregates provoke an immune response. In particular, understanding how the strength and the nature of the immune response depends on the biophysical properties of the oligomers/aggregates could be useful information. Such information could help formulate a protein-based vaccine by increasing its immunogenicity in a controlled manner [12,13].

We previously showed that ED3 of dengue serotype 3 (D3ED3) tagged with a hydrophobic 4-Ile tag generates oligomers eliciting a strong immune response with T-cell immunological memory in mouse models [16]. This study characterizes the biophysical and biochemical nature of D3ED3 oligomerized using five distinct methods and investigates their immunogenicity. The oligomers were prepared by miss-shuffling ED3’s disulfide bonds (hereafter Ms) and by using three types of stresses: Heat (Ht), Stir (St), and Freeze-thaw (FT). In addition, we used the 5-Ile tagged D3ED3 as a control **(Figure 1A and B)**. All of the D3ED3 oligomers showed a hydrodynamic radius (*R*h) of less than 100 nm (mostly between 30 around 55 nm) but had distinct biophysical (secondary and tertiary structural content) and biochemical (SS bond pattern and stability to proteolytic digestion) properties. *In vivo* mice immunization experiments showed that the Ms formulation of D3ED3, which holds a molten globular-like structure, exhibited the strongest IgG response against the native D3ED3, with long-term serum IgG level and central and effector T-cell immunological memory, as suggested by cell surface CD marker analysis. Altogether, our results suggest that controlled protein oligomerization by reshuffling the natively formed disulfide bonds could be a versatile approach for increasing a protein’s immunogenicity, and as such, it may pave the way for the development of protein-based vaccines.

**Figure 1:**
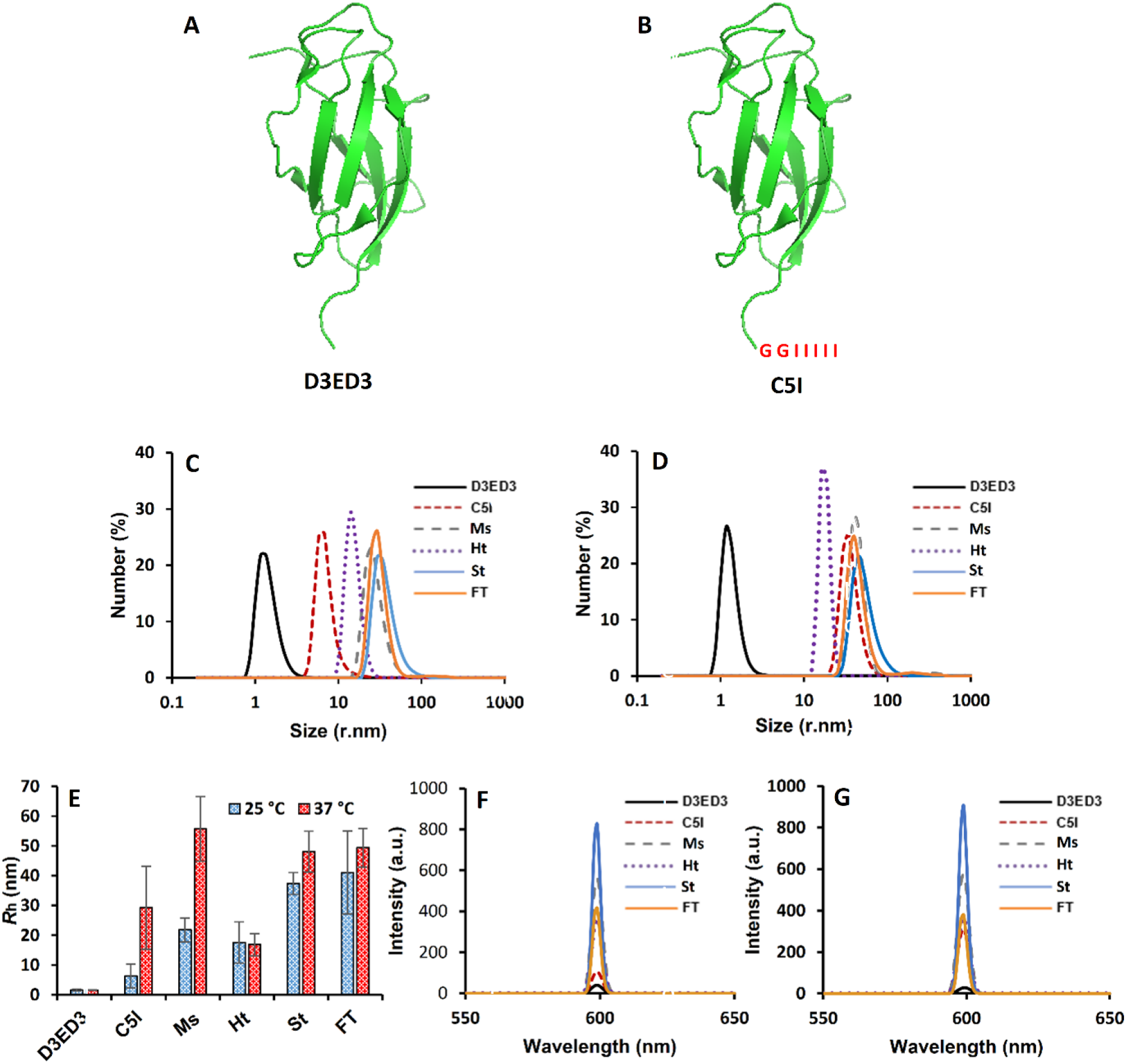
Oligomer size by dynamic light scattering (DLS) and static light scattering (SLS): **(A)** Ribbon model of D3ED3 (PDB: 3vtt) **(B)** D3ED3 with C-terminal 5 Isoleucine Tag (C5I). Hydrodynamic radii (*R*h) of D3ED3 oligomer at 0.3 mg/mL concentration measured by dynamic light scattering at **(C)** 25 °C and **(D)** 37 °C. **(E)** Temperature-dependent *R*h measured by DLS. Light scattering intensities were measured by SLS at 600 nm and at **(F)** 25 °C and **(G)** 37 °C. Line symbols are explained within the panels.

## Material and Methods

### Protein expression and purification

The D3ED3 variants were overexpressed in *E. coli* JM109(DE3) pLysS as inclusion bodies as reported earlier [16]. Protein expression was induced by the addition of 1.0 mM IPTG when the optical density (OD) of the culture medium reached 0.6 at 590 nm. After harvesting, the cells were lysed in lysis buffer (150 mM NaCl, 0.5 % sodium deoxycholate, and 1 % SDS in 50 mM Tris–HCl, pH 8.7) and lysis wash buffer (lysis buffer supplemented with 1 % v/v NP-40), and the cell lysates were air oxidized for 36 h at 30 °C in 6 M guanidine hydrochloride in 50 mM Tris–HCl, pH 8.7. The his-tagged D3ED3s were purified by Ni-NTA (Wako, Japan) chromatography, followed by dialysis against 10 mM Tris–HCl, pH 8.0 at 4 °C. The N-terminal his-tag was cleaved by thrombin proteolysis, and D3ED3 was further purified by a second round of Ni-NTA chromatography, followed by reversed-phase HPLC (Intrada 5WP-RP, IMTAK, Japan). Protein purities and identities were confirmed by analytical reversed-phase HPLC and MALDI-TOF mass spectrometry (Autoflex III, Bruker Daltonics, USA), respectively, and stored as a lyophilized powder at −30 °C until use.

### Preparation of oligomeric D3ED3

Oligomeric D3ED3s were prepared from HPLC purified native monomeric D3ED3. The lyophilized 3ED3 was dissolved in Milli-Q water (MQ) at a 1 mg/mL concentration and centrifuged at *20,000g* for 20 min at 4 °C. Samples were prepared by adding PBS to the stock solution (pH 7.4) at a 0.3 mg/mL final concentration. The heat-stressed (Ht) sample was prepared by incubating at 70 °C for 60 min using a dry heat block (MyBL-100C, AS ONE, Japan). The stir-stressed sample (St) was prepared by diluting the protein stock in MQ at a concentration of 0.6 mg/mL, and 400 μL of the protein sample was shaken at 2, 800 rpm for 75 min using a present mixer (TAITEC, Japan). Final concentrations were adjusted to 0.3 mg/mL in 1x PBS by adding MQ and 5X PBS as a stock buffer. The 0.6 mg/mL solution was also used for preparing the freeze-thaw oligomers (FT). Eight repetitive cycles of liquid nitrogen and 37 °C incubation were performed, and finally, MQ and 5x PBS were added to the sample to adjust the final concentration of the Ft sample at 0.3 mg/mL in 1x PBS.

The SS-bonds miss-shuffled oligomer (Ms) was prepared by dissolving the HPLC purified D3ED3 in 6 M guanidine hydrochloride and 10 mM Tris-HCl (pH 8.7) at a final concentration of 10 mg/mL. 200 mM of dithiothreitol (DTT; Wako, Japan) was added to the sample to break the disulfide bonds, and the sample was incubated for 3 hours at room temperature. 10 % acetic acid was added to stop the reaction, and the sample was dialyzed overnight against MQ at 4 °C. The protein sample was then again dissolved in 10 mM Tris-HCl (pH 8.7) containing 6 M guanidine hydrochloride and incubated for 6 hours at room temperature to form miss-shuffled SS bonds by air-oxidization. After 6 hours, 10 % acetic acid was added and dialyzed again against MQ at 4 °C overnight. Finally, after centrifugation (20,000 xg for 20 min), the supernatant containing soluble D3ED3 was stored at −30 °C as the Ms stock solution. The final concentration was adjusted by adding MQ and 5x PBS to a final concentration of 0.3 mg/mL in 1x PBS.

### Dynamic light scattering (DLS) and static light scattering (SLS)

The hydrodynamic radii (*R*_h_) of oligomeric D3ED3 were measured using dynamic light scattering (DLS) on a Zetasizer Nano-S (Malvern, UK) at a protein concentration of 0.3 mg/mL in PBS (pH 7.4) at 25 °C and 37 °C. 100 μL of protein sample in a polystyrene cuvette (BRAND, Germany) was used. For each sample, three individual measurements were performed and averaged to calculate the final *R*_h_ value from the number distributions.

Static light scattering (SLS) was measured under the same sample condition at a wavelength of 600 nm and temperatures of 25°C and 37 °C with an FP-8500 spectrofluorometer (JASCO, Japan). 200 μL of protein samples in a 3-mm optical path length quartz cuvette (T-507, TOSOH, Japan) was used for the measurements. The final spectra were obtained by subtracting the corresponding buffer spectrum.

### Spectroscopic measurements

All spectroscopic measurements were carried out with protein at a 0.3 mg/mL concentration in PBS (pH 7.4). Secondary structure contents were measured by far-UV circular dichroism (cd) spectroscopy using a JASCO J820 cd spectropolarimeter (JASCO, Japan). 500 μL of protein solution was placed in a 2 mm path-length quartz cuvette (TOSOH, Japan), and spectra were collected in a continuous scanning mode from 260 to 200 nm wavelength. Blank spectra were measured with 1xPBS and subtracted from the spectra.

Tryptophan and 8 anilino-1-naphthalene-sulfonic acid (ANS) fluorescence were measured on a JASCO J-820 spectropolarimeter (JASCO, Japan). Protein samples were prepared in the same way as for cd measurements at a 0.3 mg/mL concentration. ANS fluorescence measurements (SIGMA-ALDRICH, Germany) were added to the solution at a final concentration of 20 μM and incubated at room temperature for 5 min.

### Limited proteolysis

Limited proteolysis was performed using trypsin (Nacalai Tesque, Japan). D3DE3 protein samples were prepared at a concentration 0.3 mg/mL (25 μM) in PBS (pH 7.4) and mixed with trypsin at a final concentration of 1,3 and 5 μg/mL. The mixture was then incubated for 0 min, 30 min, 60 min, and 120 min at 37°C. 20 μL of the reaction mixture was sampled after an incubation period, and enzymatic proteolysis was stopped by mixing the protein sample with sodium dodecyl sulfate-polyacrylamide gel electrophoresis (SDS-PAGE) sample buffer supplemented with β–mercaptoethanol, and heating at 95 °C for 5 min. The proteolysis was further monitored by running the SDS-PAGE gel.

### High-performance size exclusion chromatography (HP-SEC)

The formation of intermolecular SS bonds was investigated by high-performance size exclusion chromatography (HP-SEC) using Shimadzu LC-20AD HPLC equipment (SHIMADZU CORP, Japan). All D3ED3 samples were prepared at a concentration of 0.1 mg/mL in PBS (pH 7.4) containing 6 M guanidine hydrochloride. The reduced state was prepared by adding 0.5 M dithiothreitol (DTT) for cleaving the disulfide bond. Before analysis, the samples were centrifuged (20,000 *g,* 20 min, 4 °C) and filtered with a 0.2 μm filter. 20 μL of the protein sample was loaded onto a TSK Gel 2000 SWXL column (Tosoh Bioscience, Japan) using PBS (pH 7.4) containing 6 M guanidine hydrochloride as a mobile phase at a flow of 0.5 mL/min. The protein was detected by a fluorescence detector (RF-20A, SHIMADZU CORP, Japan) using an excitation wavelength of 295 nm and an emission wavelength of 345 nm for tryptophan. For experiments under reduced conditions, the mobile phase was supplemented with 10 mM of DTT.

### Immunization experiments

Four-week-old female mice (Jcl:ICR, CLEA, Japan) were used for immunization experiments. In total, 20 mice were divided into seven groups (*n*=2~4). Control mice were injected with PBS (pH 7.4) only. Oligomeric D3ED3 formulations were prepared at a final concentration of 0.3 mg/mL in PBS and confirmed the oligomer size by measuring the *R*h using DLS just before each round of immunization. Mice were immunized by subcutaneous injection of 100 μL sample once a week for up to 5 weeks. Serum anti-D3ED3 antibody titers (IgG) were measured four days after each round of inoculation using tail-bleed sera by enzyme-linked immunosorbent assay (ELISA). All experiments were performed in compliance with the Tokyo University of Agriculture and Technology (TUAT) and Japanese governmental regulations on animal experimentation.

### ELISA

ELISA was carried out as previously described [17]. In short, anti-D3ED3 IgG levels were evaluated using native D3ED3 (2.5 μg/mL in PBS) as coating antigens on 96-well microtiter plates (TPP microtiter plates, Switzerland). Anti-D3ED3 sera were applied to the PBS-washed wells at an initial dilution of 1:50 for tail-bleed samples. Plates were then incubated at 37 °C for 2 hours. After washing the plates thrice with PBS-0.05% Tween-20, each well was filled with 100 μL of anti-mouse IgG HRP conjugate (Thermo Fisher Scientific, USA) at a 1:3000 dilution in 0.1 % BSA-PBS-Tween-20 and incubated at 37 °C for 1.5 hours. As a substrate, O-phenyl Di-amine (OPD; TCI, Japan) was added. Color intensities were measured at 492 nm using a microplate reader (SH-9000 Lab, Hitachi High-Tech Science, Japan) immediately after stopping the reaction with 1 N sulfuric acid, 50 μL/well (Wako, Japan). Finally, titers were calculated using a power fitting model, and values were averaged over the number of mice (*n*) in the respective groups.

### Flow cytometry

Flow cytometry samples were prepared according to our previously reported protocol [16,19]. In short, single-cell suspension of mice splenocytes was prepared in FACS buffer (PBS supplemented with 2 % FBS, 1 mM EDTA, and 0.1 % sodium azide). Afterward, 1X red blood cell (RBC) lysis buffer (0.15 M ammonium chloride, 10 mM potassium bicarbonate, 0.1 mM EDTA) was used to lysis the RBCs. Further, 1 million splenocyte cells in 100 μL of pre-cooled FACS buffer were surface stained with fluorescence labelled antibodies according to the manufacturer’s guidelines. The CD4 T-lymphocytes were stained with anti-CD3-Pcy5, CD4-Pcy7, CD44-FITC, and CD62L-PE-conjugated antibodies in one tube, and CD8 T-lymphocytes were stained with anti-CD3-Pcy5, CD8-Pcy7, CD44-FITC, and CD62L-PE-conjugated antibodies (BioLegend, United State) in another tube (0.2 μg of antibodies/100 μL) for 30 min in the dark. Unbound conjugated antibodies were removed by centrifugation. Finally, cells were resuspended in 1 mL of FACS buffer, and the data were collected using CytoFlex (Beckman Coulter, USA).

## Results

### Biophysical and biochemical properties

#### D3ED3 Oligomeric Sizes

We determined the oligomeric size of D3ED3 by measuring the hydrodynamic radius (*R_h_*) using dynamic light scattering (DLS). Measurements were performed under conditions identical or nearly identical to those used for mice immunization (0.3 mg/mL in PBS, pH 7.4) at 25 °C and 37 °C. At both temperatures, the *R*h of the unstressed and untagged D3ED3 was around 1.78 nm ± 0.39 nm **(Figure 1C-E and Table 1)**, as expected for a small monomeric globular protein with a molecular weight of ~11 kDa and in line with our previous observations [15]. The disulfide bonds miss-shuffled oligomer (Ms), stir (St), and freeze-thaw (FT) formulation of D3ED3 showed an *R*h of approximately 50 nm at 37 °C, indicating their oligomeric state **(Figure 1E and Table 1)**. C5I oligomerized in a temperature-dependent manner, and *R*h increased from 6 nm to ~30 nm when the temperature was raised from 25 °C to 37 °C. No reversibility was observed, unlike for BPTI tagged with Isoleucine (BPTI-C5I) [20]. The probable mechanism of oligomer formation through C-terminal Isoleucine was discussed in our previous studies with BPTI [21]. The heat-induced D3ED3 (Ht) produced the smallest oligomer with an *R*h of 28.15 ± 15.75 nm.

**Table 1.** Oligomeric status of D3ED3 formulations by DLS and SLS at 37 °C

| D3ED3 formulation | DLS | SLS |
| --- | --- | --- |
|  | (*R<sub>h</sub>*, nm) | (Intensity, a.u.) |
| D3ED3 | 1.78 ± 0.39 | 40.46 |
| C5I_25 °C | 6.29 ± 3.99 | 104.51 |
| C5I_37 °C | 29.26 ± 5.56 | 310.51 |
| Ms | 55.73 ± 10.76 | 561.17 |
| Ht | 28.15 ± 15.75 | 362.06 |
| St | 48.03 ± 6.89 | 825.64 |
| FT | 49.44 ± 6.47 | 416.53 |

The degree of oligomerization was further characterized by static light scattering (SLS). The light scattering intensity of monomeric D3ED3 was minimal. In contrast, all stress-induced D3ED3 (Ht, St, and FT), C5I, and Ms showed a strong light scattering both at 25 °C and 37 °C, confirming their oligomerization and corroborating with DLS data **(Figure 1F, G, and Table 1)**.

#### Secondary structure content and tertiary structure stability

The secondary structure content of St and FT D3ED3 at 37 °C, as characterized by far UV-cd (200-260 nm), was essentially identical to that of the native monomeric D3ED3 **(Figure 2A)**. Heat incubation at 70 °C for 60 min (Ht) significantly altered the cd spectrum of D3ED3 **(Figure 2A)**, which might indicate a decrease of antiparallel β-sheet strands according to BeStSel (**Table S1)**. The spectrum of the Ms was intermediate between the monomeric D3ED3 and the heat-induced D3ED3 oligomers (Ht). The secondary structure content as assessed by cd value at 220 nm was thus ranked as follows: Ht> C5I> Ms> St~ FT~monomeric D3ED3.

**Figure 2:**
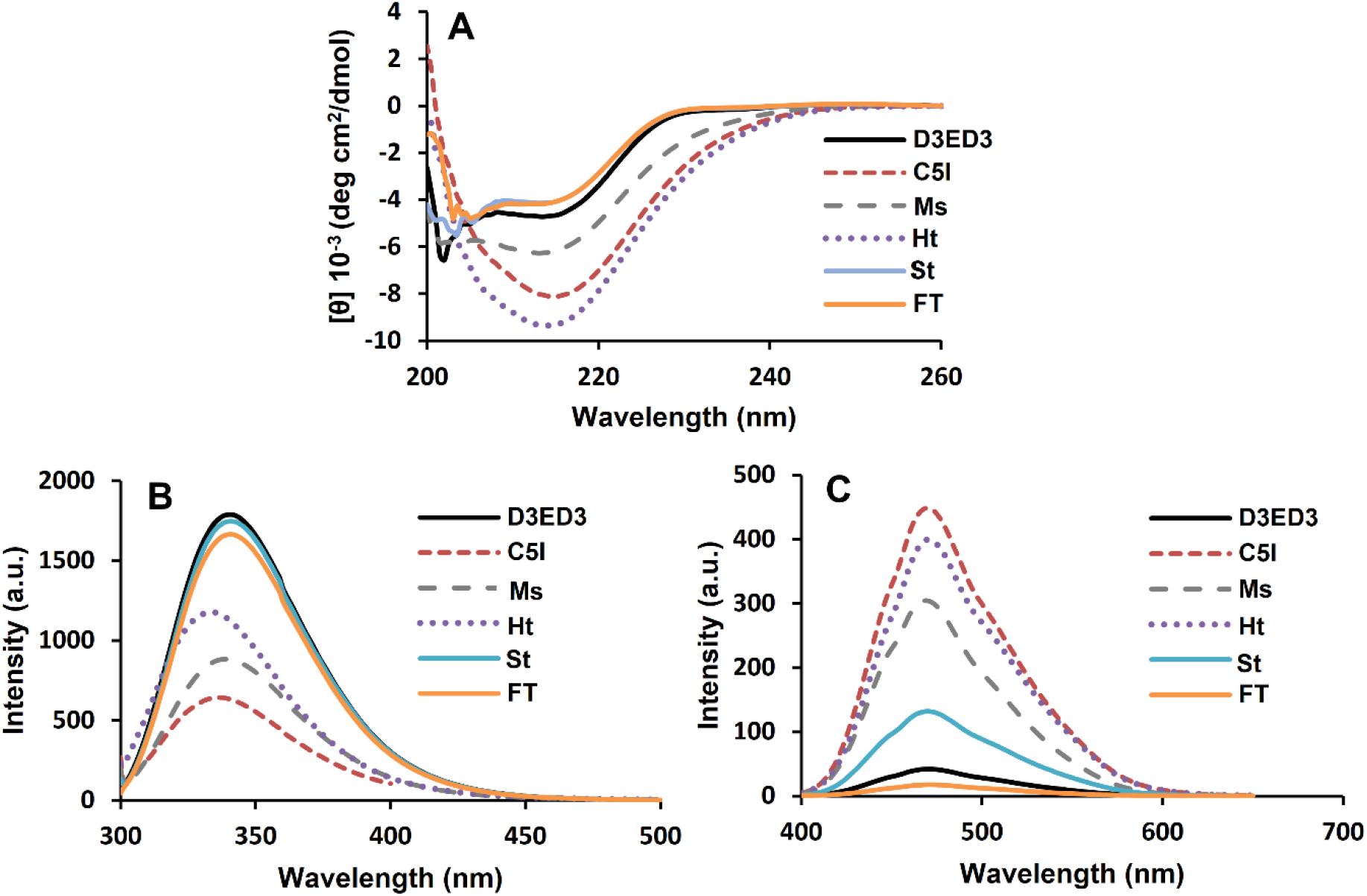
Secondary and Tertiary Structure characterization: **(A)** Far-UV circular dichroism spectrum measured at a protein concentration of 0.3 mg/m at 37 °C. Tryptophan **(B)** and ANS-fluorescence intensity **(C)** measured under the same conditions as for **A**. Line symbols are explained within the panels.

The native-likeness of the tertiary structure was characterized based on tryptophan and ANS fluorescence. Fluorescence intensity is highly sensitive to the microenvironment surrounding the tryptophan sidechains and provides valuable information on protein interiors’ rigidity and loose packing [24]. The monomeric D3ED3 showed a strong tryptophan fluorescence as expected for a natively packed protein **(Figure 2B)**. The tryptophan fluorescence intensity of St and FT was essentially identical to the monomeric D3ED3, which was in line with the observation of unaltered secondary structure measured by cd, suggesting that in St and FT D3ED3 retains a native-like structure in the oligomeric state. On the other hand, the fluorescence emission intensity of C5I, Ms, and Ht was significantly weaker than that of the native D3DE3, suggesting a molten-globule-like, loosely packed interior enabling water molecules to access the tryptophan side chain and quench fluorescence [23]. The tryptophan fluorescence ranked as: monomeric D3ED3~St~FT>Ht>Ms>C5I.

Next, we used ANS, which is indicative of a molten-globule state and emits fluorescent light upon binding to partially exposed and loosely packed hydrophobic pockets [30]. The monomeric D3ED3 showed the lowest ANS fluorescence intensity, as expected for a compact, native structure **(Figure 2C)**. On the other hand, C5I and Ht exhibited the strongest ANS fluorescence intensity, confirming the molten-globule-like state as indicated by tryptophan fluorescence [24]. FT’s ANS-fluorescence intensity was very close to that of the monomeric D3ED3, whereas the fluorescence intensity of St increased marginally. Ms’ fluorescence was significantly stronger than that of the native monomeric D3ED3 but slightly weaker than C5I and Ht. Overall, the ANS fluorescence intensity ranked as follows: C5I> Ht> Ms> St> FT~monomeric D3ED3.

Further, a limited proteolysis experiment was performed to check the biochemical stability of the oligomeric D3ED3. Monomeric D3ED3, St, and FT showed very similar stability, whereas the Ht sample showed minimum stability against trypsin digestion **(Figure 3)**. Ms showed stability higher than that of Ht but lower than D3ED3, in line with their intermediate secondary structure content and tertiary structure stability measured by cd and fluorescence spectroscopy, respectively.

**Figure 3:**
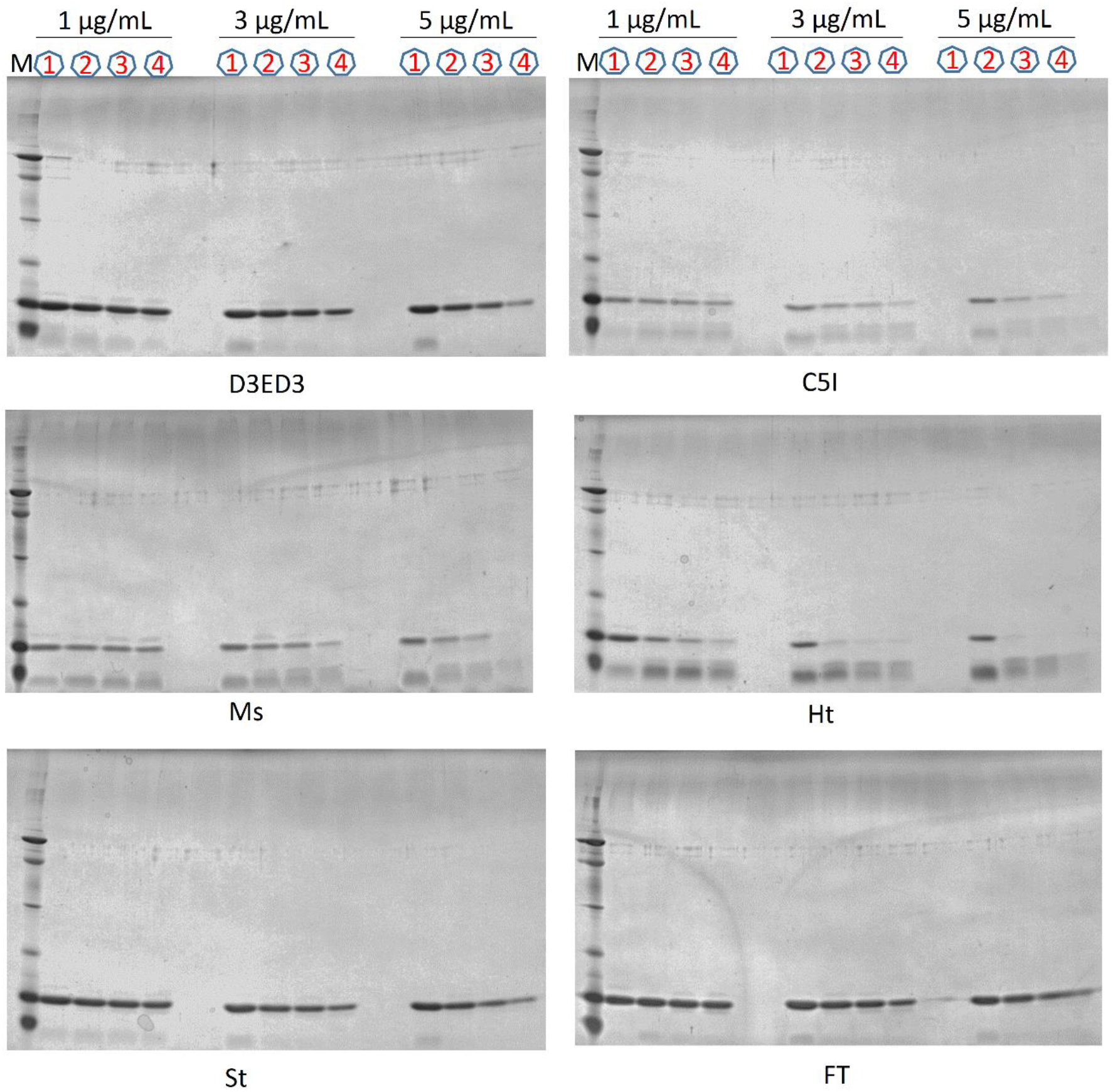
Limited proteolysis. SDS-PAGE gel showing the trypsin digestion of oligomeric D3ED3. Circled 1, 2, 3, and 4 indicate the 0, 30, 60, and 120 min incubation sample bands, respectively.

#### Characterization of disulfide bond pattern

We further characterized the presence of intermolecular SS bonds using high-performance size exclusion chromatography (HP-SEC). HP-SEC Samples were performed in the presence of denaturant to decrease intermolecular interaction. D3ED3, C5I, Ht, St, and FT showed a single elution peak under reducing and oxidizing conditions. The elution time was around 15 min, corresponding to the molecular weight of the monomeric D3ED3 and indicating the absence of intermolecular SS bonds **(Figures 4A and B)**. On the other hand, Ms showed a smaller monomeric peak under oxidizing conditions, and a broad peak at a higher molecular weight appeared. The high molecular weight peak disappeared under reducing conditions, indicating the presence of intermolecular SS bonds in the Ms sample **(Figures 4A and B)**.

**Figure 4:**
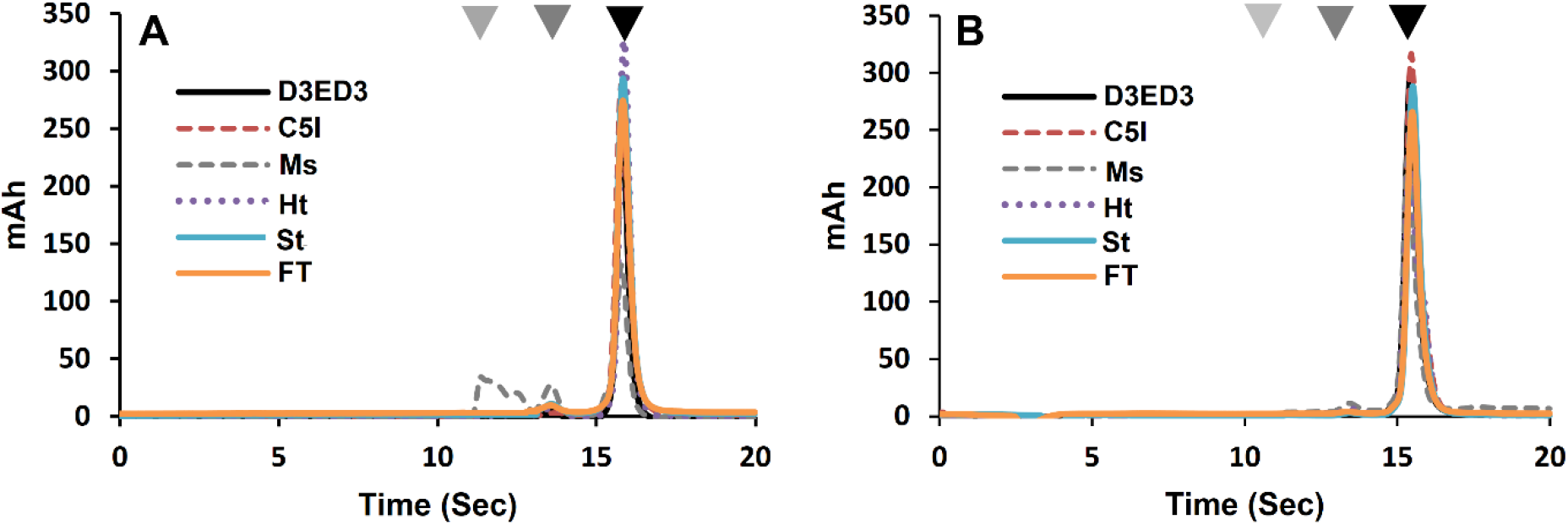
Characterization of intermolecular SS bond: The presence of intermolecular SS bond was determined by high-performance size exclusion chromatography (HP-SEC). 20 μL of denatured D3ED3 sample was used for HP-SEC analysis in a non-reducing (A) and a reducing (B) condition. Gray, dark gray and black triangles indicate the trimer, dimer, and monomer D3ED3 elusion peak, respectively, and line symbols are explained within the panels.

### Effect of D3ED3 oligomerization type on immune response

The *in vivo* immunogenicity of D3ED3 was evaluated using Jcl:ICR mice. As in all our studies, we confirmed the size of the oligomer just before injection into the mice because the aggregates might be sensitive to small changes in external conditions and might form accidentally **(Supplementary Figure S1)**. The monomeric D3ED3 was scarcely immunogenic as assessed by serum anti-D3ED3 IgG titer using ELISA **(Figure 5A)**. This is in line with our previous reports [15] and corroborates that small proteins are not or are poorly immunogenic [31]. The C5I tagged D3ED3 showed an increased serum anti-D3ED3 IgG titer than monomeric D3ED3, whereas Ms showed the strongest IgG titer **(Figure 5A)**. The immune response of Ht, St, and FT was weak, although those formulations, particularly St and FT contain oligomer’s size very similar to Ms **(Figure 1C and Table 1)**. IgG subclass determination by ELISA revealed that all of the mice (3 out of 3) immunized with Ms developed IgG1 type immune response **(Figure 5C)**.

**Figure 5:**
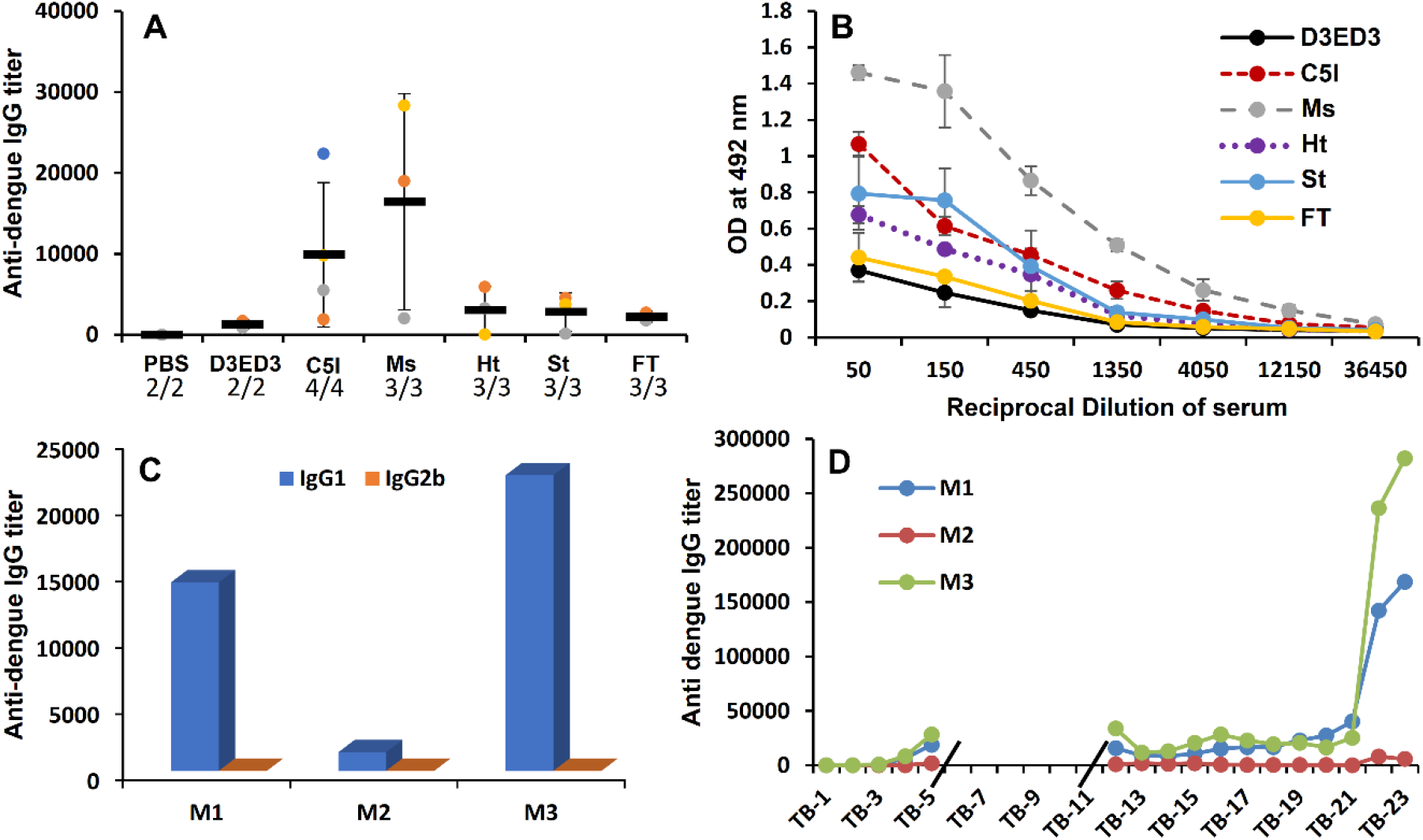
Anti-D3ED3 IgG response by ELISA and *in vivo* immunological memory: ELISA was performed using tail bleed (TB) serum samples after five doses of subcutaneous immunization. ELISA plates were coated with the native monomeric D3ED3. **(A)** anti-D3ED3 IgG titer of different D3ED3 formulations. The dots stand for the titer of the individual mice. All mice titers are shown, including non-responsive mice. The average is taken on the responsive mice only. **(B)** Absorbance at 492 nm versus the reciprocal dilution of anti-sera developed against D3ED3 formulations. **(C)** IgG subclass of Ms immunized mice. **(D)** Long-term anti-D3ED3 serum IgG titer of Ms immunized mice and IgG titer after re-immunization (boost shot) with the native monomeric D3ED3.

### Ms generated immunological memory

In order to assess the persistency of the immune response, we kept the Ms immunized mice for 17 weeks after the last injection (5^th^ round). We observed a long-lasting serum anti-D3ED3 IgG **(Figure 5D)**. Furthermore, a booster shot using the native monomeric D3ED3 injected into the Ms-immunized mice twenty-two weeks after the first injection elicited a rapid and sharp increase in the IgG titer, which was 100 times stronger than the primary immunization **(Figure 5D)**. Thus, immunization with Ms generated immunological memory against the native D3ED3.

Splenocytes collected 23 weeks after the first immunization were analyzed by cell surface cluster of differentiation (CD) marker. Mice immunized with monomeric D3ED3 had a low percentage of CD44+ and a high percentage of CD62L+ CD4 (helper T-cell) and CD8 (cytolytic T-cell) cell populations **(Supplementary Figure S2)**. Moreover, a higher number of CD44-CD62L+ co-expressed cell populations indicated their naïve immunological status **(Figure 6A and B)** [27,28]. On the other hand, Ms-immunized mice showed a higher CD44+ cell population **(Supplementary Figure S2)**. Additionally, we observed a higher percentage of CD4 and CD8 T-cell co-expressing CD44+CD62+ and CD44+CD62L- **(Figures 6A and B)**, strongly suggesting the generation of central and effector T-cell memory [29].

**Figure 6:**
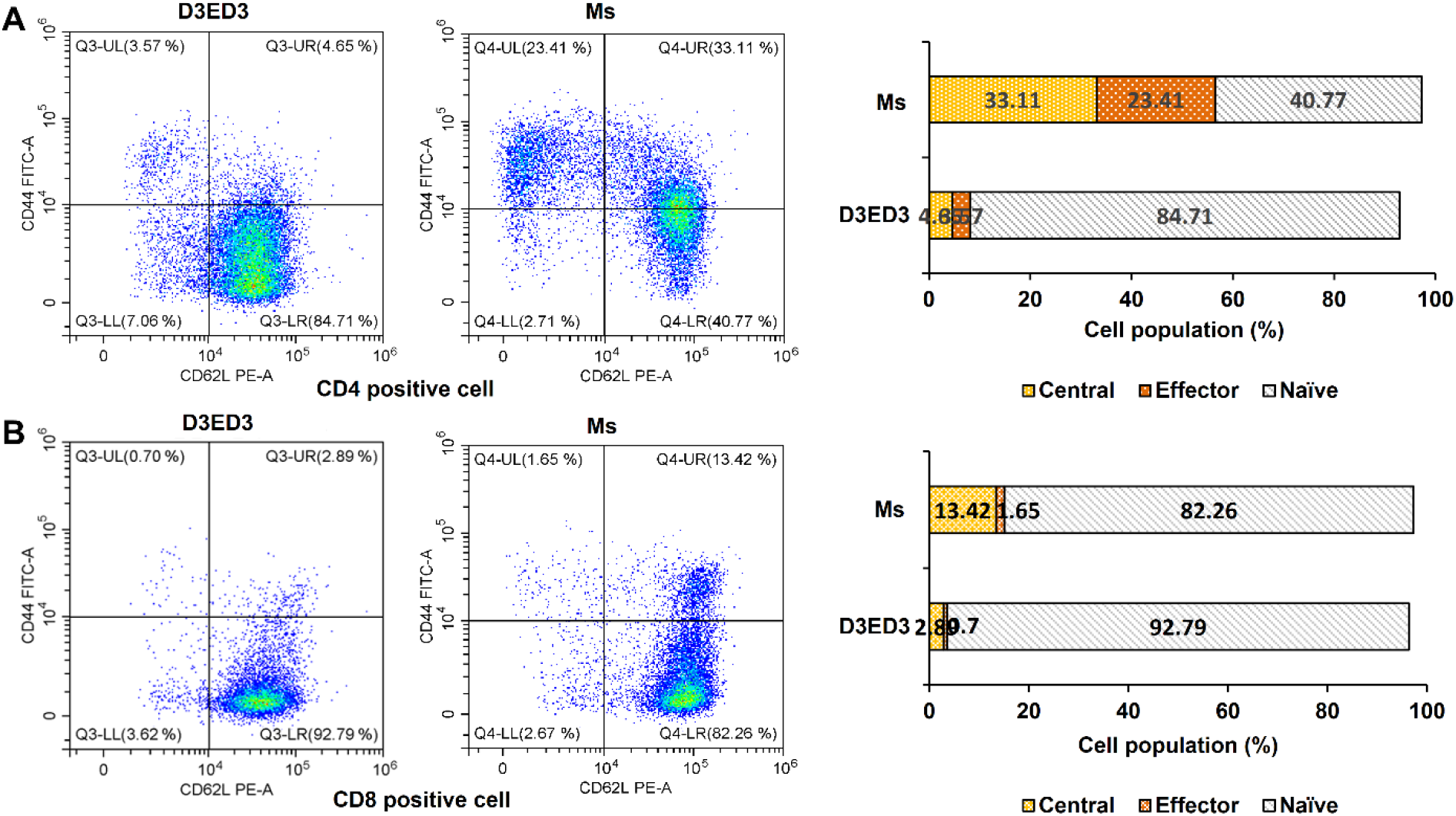
Flow cytometry analysis of cell surface CD maker: Immunological memory was characterized by analyzing the cell surface CD marker. A single-cell suspension of mice splenocytes in FACS buffer was used for flow cytometry. Differential expression of surface CD marker **(A)** on CD4+ cells (T-helper) and **(B)** on CD8+ cells (T-cytolytic).

## Discussion

Protein oligomerization has been reported to increase immunogenicity, which is usually rationalized by the presence of multiple repetitive B-cell epitopes formed through oligomerization [8]. However, the relationship between oligomerization (or aggregation) and immunogenicity remains a matter of some debate [29–31]. For example, oligomers of natively folded structural factor VIII are immunogenic [32], whereas the oligomers of fully unfolded or denatured factor VIII are not or little immunogenic [33]. On the other hand, oligomers of natively folded functional mAbs (according to dot blotting) generated by freeze-thaw or shaking are not immunogenic, but they can turn immunogenic upon chemical modification, such as oxidation [34].

Our study showed that the way amorphous oligomers are produced would determine their biophysical and biochemical nature, which in turn affected the immunogenicity of the proteins. Further, our observation indicates that the amorphous oligomers **(Supplementary Figure S3)** formed by disulfide bond scrambling and in a molten globular-like state had the strongest tendency to increase the IgG response. On the other hand, the other oligomers, even though having almost identical sizes, were not immunogenic, supporting our previous report performed with V_HH_ [15]. In addition, the D3ED3 oligomer formed through disulfide bond scrambling was even more potent than the C5I-induced oligomers we reported previously **(Figure 5A)** [16]. Moreover, a booster shot with native monomeric D3ED3 elicited a strong anti-D3ED3 IgG response **(Figure 5D)**, suggesting the establishment of immunological memory, which is required from a vaccine candidate [35].

CD marker analysis indicated the induction of central and effector T-cell memory **(Figure 5)**. The generation of T-cell response begins with the uptake of antigen by professional antigen-presenting cells (APCs) and their subsequent migration into draining lymph nodes, where T-cell responses are initiated in parallel with B-cell response [36]. We thus speculate that the strong immunogenic response initiated by Ms **(Figure 5A)** might be related to a higher antigen uptake by APCs in the oligomeric than in the monomeric state [10].

### Conclusions

Antibodies directed against envelope protein domain 3 (ED3) are effective in reducing the viral load and could provide protective immunity against dengue infection [37]. However, dengue ED3 is poorly immunogenic, and adjuvants are needed [38]. We showed that amorphous aggregates of D3ED3 oligomerized in a molten-globule-like state (C5I and Ms) elicited a robust immune response against the natively folded monomeric proteins. On the other hand, amorphous aggregates where D3ED3 are oligomerized in a native-like state (Ht, St, and FT) did not yield a significant response. This observation was against our expectations and the multiple-epitope presentation hypothesis, which has underlined the development of virus-like particles (VLP). To date, the VLP technique, where natively folded proteins are regularly arranged on a particle’s surface, has recently provided a vaccine against malaria [15]. Our study suggests that amorphous oligomers made of proteins produced in *E.coli* could be an attractive alternative for producing versatile and cost-effective subunit vaccines.

## Abbreviations

D3ED3: Dengue virus 3 envelop protein domain 3
cd: Circular Dichroism
DLS: Dynamic Light Scattering
SLS: Static Light Scattering
ELISA: Enzyme-linked Immunosorbent Assay
HP-SEC: High-Performance Size Exclusion Chromatography

## Acknowledgments

We thank all members of the Kuroda Laboratory for discussion and technical assistance. We are grateful to Professors Tsuyoshi Tanaka, Tomoko Yoshino and Atsushi Arakaki for the use of ZetaNanosizer equipment.

## Contribution

Y.K. and M.G.K. designed the project, and wrote the manuscript. S.B. discussed and advised experimental data, S.M. and M.G.K. performed the experiments and analyzed the data. All authors read and approved the manuscript.

## Data Sharing

All data are given in the manuscript and the supplementary data.

## Supporting Information

**Figure S1:**
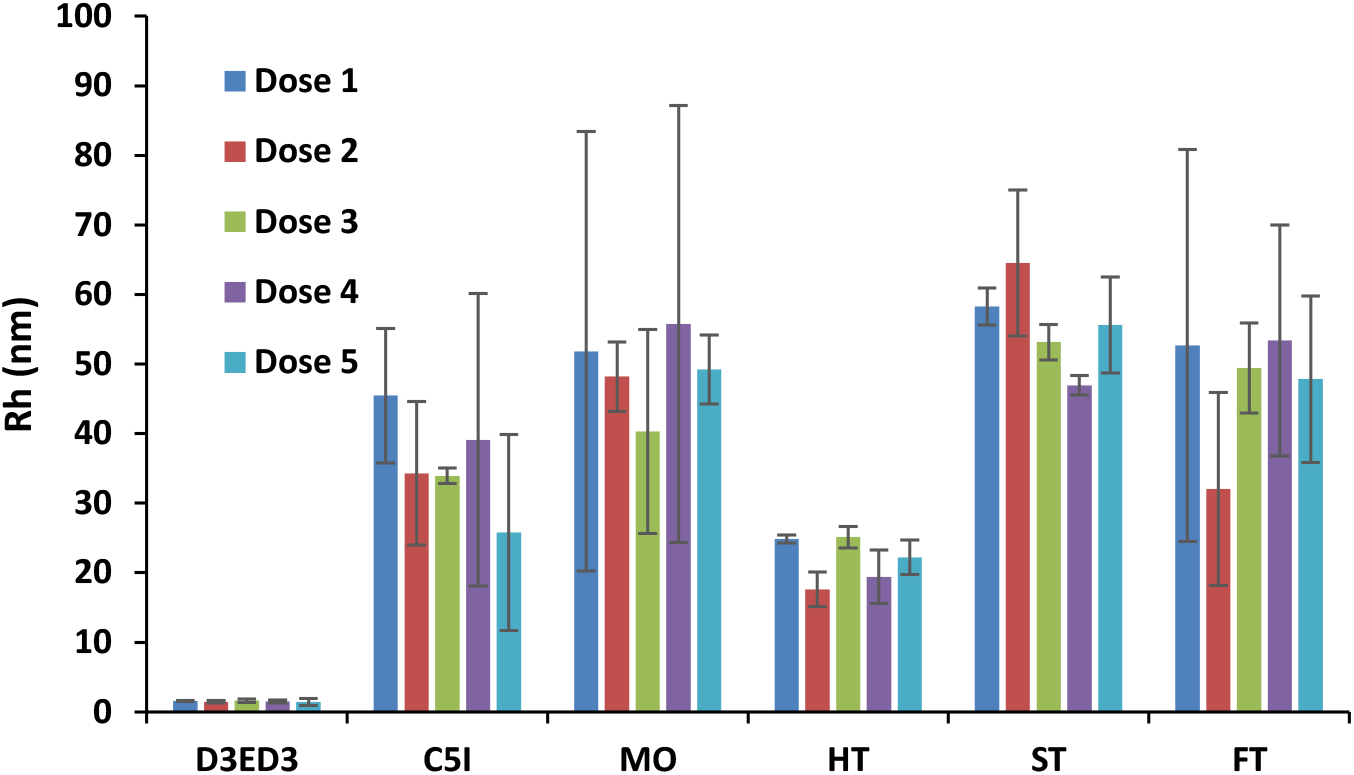
Oligomeric size measured by DLS before each round of immunization. Hydrodynamic radii (*R*h) of the D3ED3 formulations were measured just before the immunization from doses 1 to 5 at 37 °C. Line symbols are explained within the panel. Values are calculated from three independent measurements.

**Figure S2:**
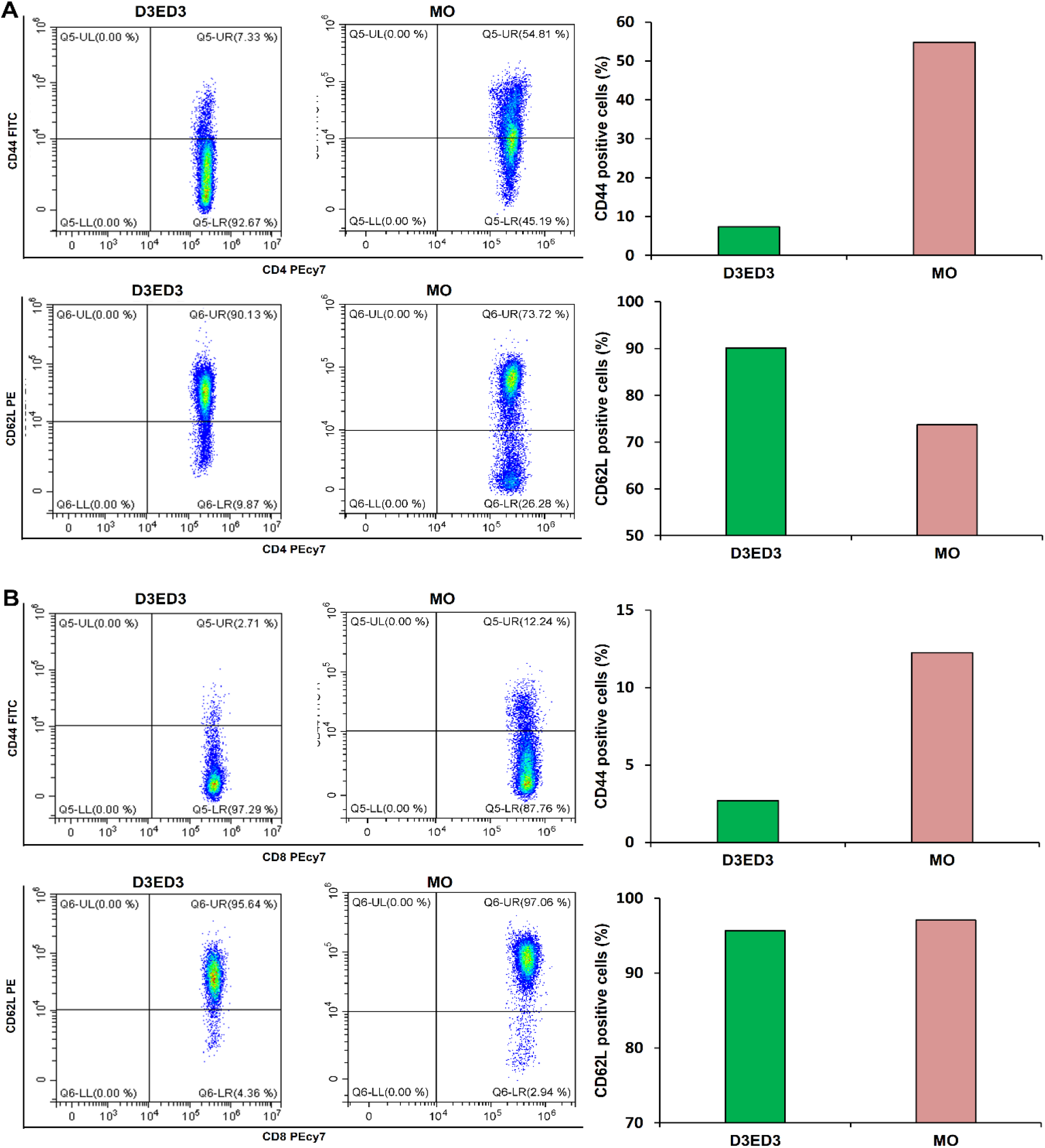
Flow-cytometry analysis of cell surface CD (cluster of differentiation) markers. A single-cell suspension of mice splenocytes in FACS buffer was used for flow-cytometry analysis. Differential expressions of cell surface CD markers induced by native D3ED3 and MO on **(A)** T-helper cell (CD4+) and **(B)** T-cytolytic cell (CD8+). MO-immunized mice had a higher population (%) of CD44+ Th-cell and Tc-cell than native D3ED3 immunized mice.

**Figure S3:**
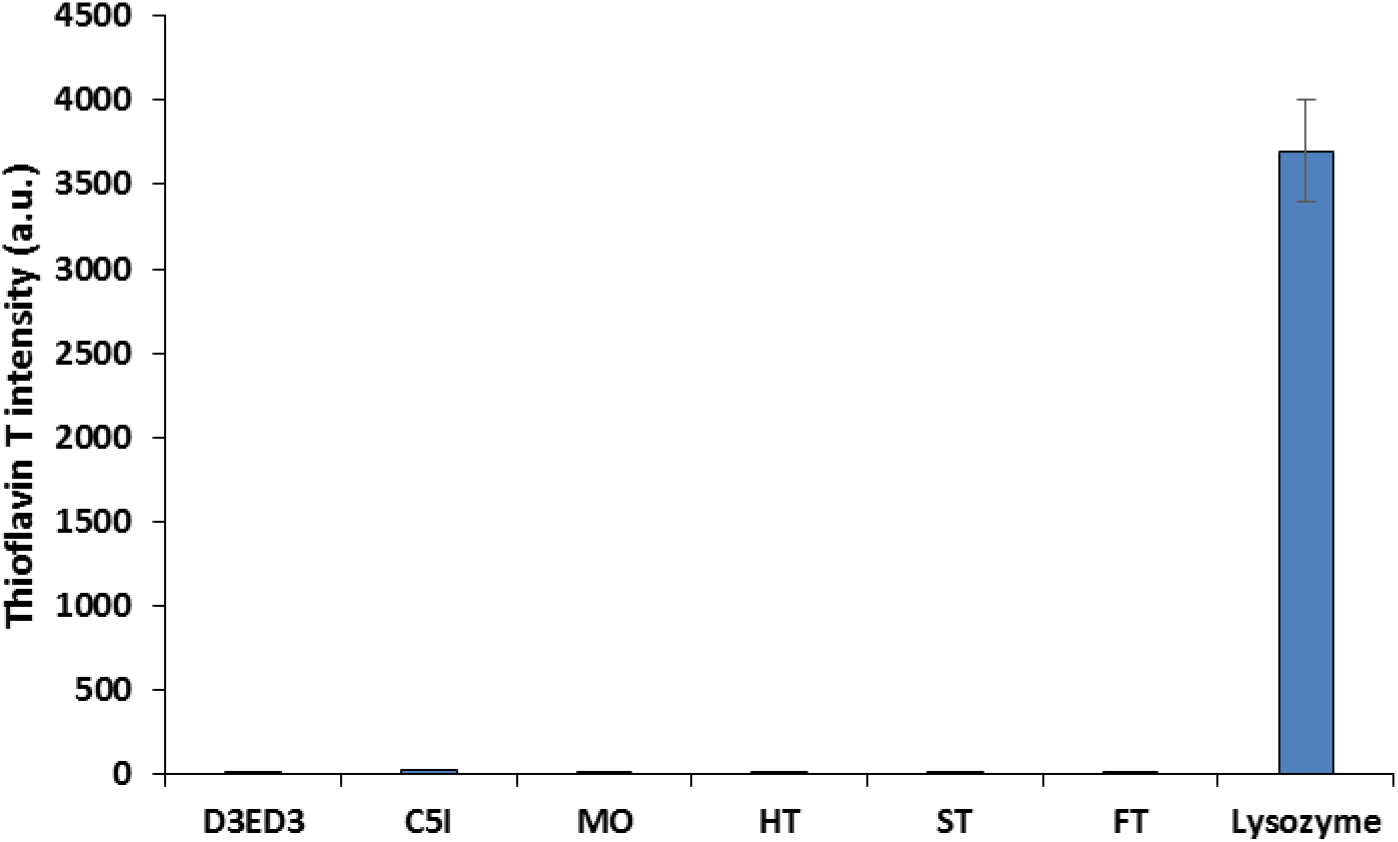
Thioflavin T assay. The presence of amyloidogenicity was assessed by Thioflavin T assay, which specifically binds to the β-sheeted structure of amyloid aggregates and gives a strong fluorescence. Lysozyme was used as a positive control for amyloidogenic aggregation. The fluorescence intensity of all of the six D3ED3 formulations was significantly lower than that of the lysozyme signal, indicating their amorphous nature.

### Measurement protocol

ED3 at a concentration of 0.3 mg/mL in PBS (pH 7.4) was used. Thioflavin T was added to the solution at a final concentration of 10 μM and incubated at 25 °C for 5 min in the dark before measurement. The fluorescence measurements were performed on an FP-8500 spectrofluorometer (JASCO, Tokyo, Japan) and using a glass cuvette containing 200 μL of the sample at 37 °C. The excitation and emission wavelength were set to 440 nm and 470 nm, respectively. Lysozyme was used as a positive control, which forms amyloid-like fibril by incubating the sample at 57 °C for 60 hours at pH 2.0 (at a protein concentration of 34 mg/mL) [1, 2].

**Table S1:**
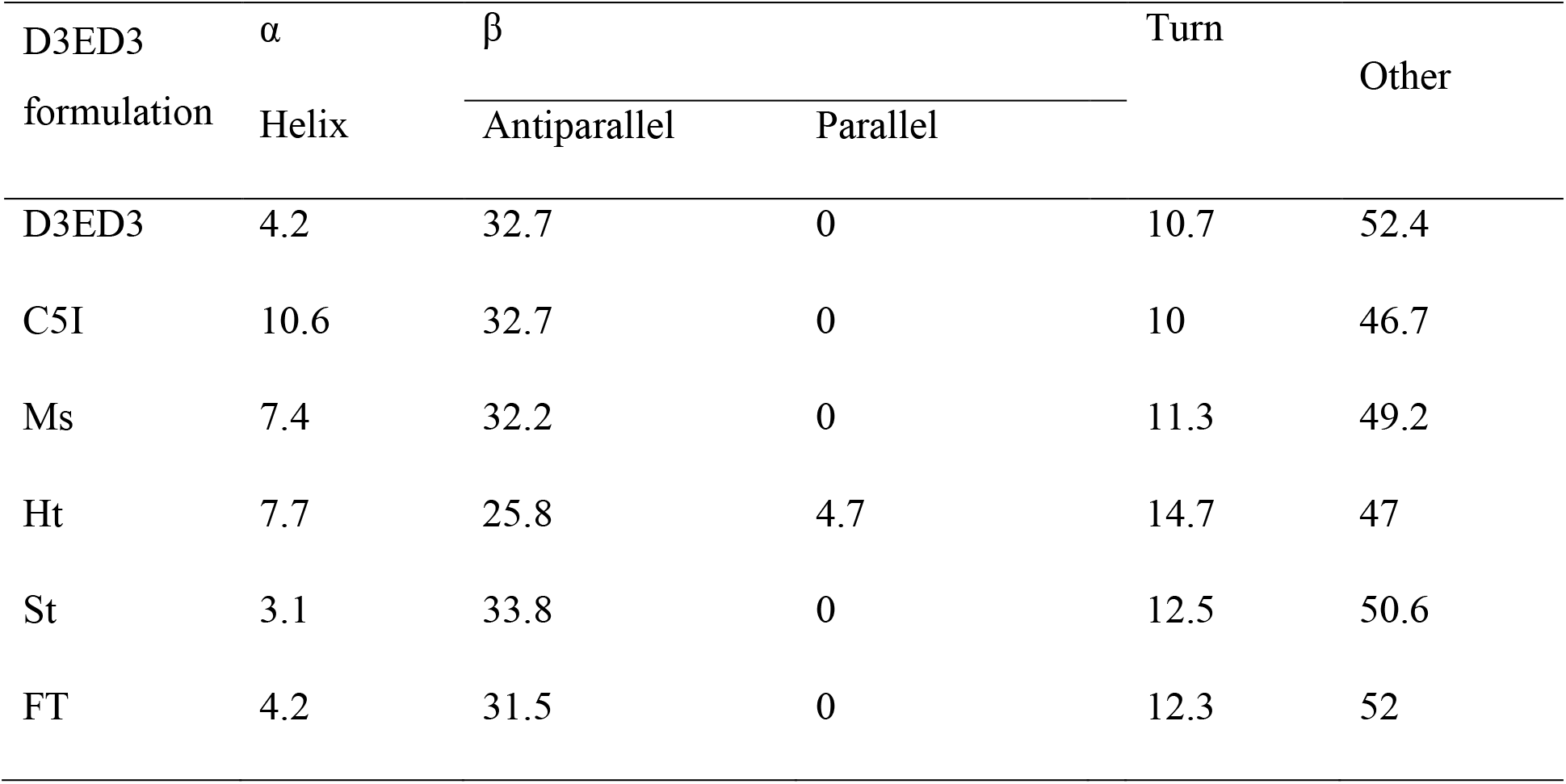
Secondary structure content (%) by BeStSel

## References

1. Chen, Y., et al., Dengue virus infectivity depends on envelope protein binding to target cell heparan sulfate. Nature Medicine, 1997. 3(8): p. 866–871.

2. Whitehead, S.S., et al., Prospects for a dengue virus vaccine. Nature Reviews Microbiology, 2007. 5(7): p. 518–528.

3. Foged, C., Subunit vaccines of the future: the need for safe, customized and optimized particulate delivery systems. Therapeutic Delivery, 2011. 2(8): p. 1057–1077.

4. Tripathi, N.K. and A. Shrivastava, Recent Developments in Recombinant Protein-Based Dengue Vaccines. Frontiers in Immunology, 2018. 9(1919).

5. Karch, C.P. and P. Burkhard, Vaccine technologies: From whole organisms to rationally designed protein assemblies. Biochem Pharmacol, 2016. 120: p. 1–14.

6. Perrie, Y., et al., Vaccine adjuvant systems: enhancing the efficacy of sub-unit protein antigens. Int J Pharm, 2008. 364(2): p. 272–80.

7. Di Pasquale, A., et al., Vaccine Adjuvants: from 1920 to 2015 and Beyond. Vaccines, 2015. 3(2): p. 320–43.

8. Moussa, E.M., et al., Immunogenicity of Therapeutic Protein Aggregates. J Pharm Sci, 2016. 105(2): p. 417–430.

9. Joshi, V.B., S.M. Geary, and A.K. Salem, Biodegradable particles as vaccine antigen delivery systems for stimulating cellular immune responses. Hum Vaccin Immunother, 2013. 9(12): p. 2584–90.

10. Nguyen, B. and NH. Tolia, Protein-based antigen presentation platforms for nanoparticle vaccines. npj Vaccines, 2021. 6(1): p. 70.

11. Wu, Y., et al., Particle-based platforms for malaria vaccines. Vaccine, 2015. 33(52): p. 7518–24.

12. Grgacic, E.V. and D.A. Anderson, Virus-like particles: passport to immune recognition. Methods, 2006. 40(1): p. 60–5.

13. Ratanji, K.D., et al., Immunogenicity of therapeutic proteins: influence of aggregation. J Immunotoxicol, 2014. 11(2): p. 99–109.

14. Rombach-Riegraf, V., et al., Aggregation of human recombinant monoclonal antibodies influences the capacity of dendritic cells to stimulate adaptive T-cell responses in vitro. PLoS One, 2014. 9(1).21.

15. Kibria, M.G., et al., The immunogenicity of an anti-EGFR single domain antibody (V(HH)) is enhanced by misfolded amorphous aggregation but not by heat-induced aggregation. Eur J Pharm Biopharm, 2020. 152: p. 164–174.

16. Islam, M.M., et al., Anti-Dengue ED3 Long-Term Immune Response With T-Cell Memory Generated Using Solubility Controlling Peptide Tags. Frontiers in Immunology, 2020. 11(333).

17. Brindha, S. and Kuroda Y. A multi-disulfide receptor-binding domain (RBD) of the Sars-CoV-2 spike protein expressed in E.coli using a sep-tag produces antisera interacting with the mammalian cell expressed spike (s1) protein. Int J Mol Sci. 2022 23(3):1703.

18. Kulkarni, M.R., et al., Structural and biophysical analysis of sero-specific immune responses using epitope grafted Dengue ED3 mutants. Biochim Biophys Acta, 2015. 10(43): p. 6.

19. Kibria, M. G., et al., Immune response with long-term memory triggered by amorphous aggregates of misfolded anti-EGFR VHH-7D12 is directed against the native VHH-7D12 as well as the framework of the analogous VHH-9G8. European Journal of Pharmaceutics and Biopharmaceutics, 2021. 165: p. 13–21.

20. Rahman, N., et al., Nanometer-Sized Aggregates Generated Using Short Solubility Controlling Peptide Tags Do Increase the In Vivo Immunogenicity of a Nonimmunogenic Protein. Mol Pharm, 2020. 17(5): p. 1629–1637.

21. Nakamura, S., et al., Reversible Oligomerization and Reverse Hydrophobic Effect Induced by Isoleucine Tags Attached at the C-Terminus of a Simplified BPTI Variant. Biochemistry, 2020. 59(39): p. 3660–3668.

22. Vivian, J.T. and P.R. Callis, Mechanisms of Tryptophan Fluorescence Shifts in Proteins. Biophysical Journal, 2001. 80(5): p. 2093–2109.

23. Kuroda, Y., S. Endo, and H. Nakamura, How A Novel Scientific Concept Was Coined the “Molten Globule State”: Biomolecules. 2020 Feb 10;10(2):269. doi: 10.3390/biom10020269.

24. Hawe, A., M. Sutter, and W. Jiskoot, Extrinsic fluorescent dyes as tools for protein characterization. Pharm Res, 2008. 25(7): p. 1487–99.

25. Rahman, N., et al., A systematic mutational analysis identifies a 5-residue proline tag that enhances the in vivo immunogenicity of a non-immunogenic model protein. FEBS Open Bio, 2020. 10(10): p. 1947–1956.

26. Bannard, O., M. Kraman, and D. Fearon, Pathways of memory CD8+ T-cell development. Eur J Immunol, 2009. 39(8): p. 2083–7.

27. Kaech, S.M., E.J. Wherry, and R. Ahmed, Effector and memory T-cell differentiation: implications for vaccine development. Nat Rev Immunol, 2002. 2(4): p. 251–62.

28. Roberts, A.D., K.H. Ely, and D.L. Woodland, Differential contributions of central and effector memory T cells to recall responses. J Exp Med, 2005. 202(1): p. 123–33.

29. Fathallah, A.M., et al., The Effect of Small Oligomeric Protein Aggregates on the Immunogenicity of Intravenous and Subcutaneous Administered Antibodies. J Pharm Sci, 2015. 104(11): p. 3691–3702.

30. Kijanka, G., et al., Submicron Size Particles of a Murine Monoclonal Antibody Are More Immunogenic Than Soluble Oligomers or Micron Size Particles Upon Subcutaneous Administration in Mice. J Pharm Sci, 2018. 107(11): p. 2847–2859.

31. Ratanji, K.D., et al., Editor’s Highlight: Subvisible Aggregates of Immunogenic Proteins Promote a Th1-Type Response. Toxicol Sci, 2016. 153(2): p. 258–70.

32. Pisal, D.S., et al., Native-Like Aggregates of Factor VIII Are Immunogenic in von Willebrand Factor Deficient and Hemophilia a Mice. Journal of Pharmaceutical Sciences, 2012. 101(6): p. 2055–2065.

33. Purohit, V.S., C.R. Middaugh, and S.V. Balasubramanian, Influence of aggregation on immunogenicity of recombinant human Factor VIII in hemophilia A mice. Journal of Pharmaceutical Sciences, 2006. 95(2): p. 358–371.

34. Filipe, V., et al., Immunogenicity of different stressed IgG monoclonal antibody formulations in immune tolerant transgenic mice. MAbs, 2012. 4(6): p. 740–52.

35. Siegrist, C.-A., 2 - Vaccine Immunology, in Plotkin’s Vaccines (Seventh Edition), S.A. Plotkin, et al., Editors. 2018, Elsevier. p. 16–34.e7.

36. Randolph, G.J., V. Angeli, and M.A. Swartz, Dendritic-cell trafficking to lymph nodes through lymphatic vessels. Nat Rev Immunol, 2005. 5(8): p. 617–28.

37. Chávez, J.H., et al., Domain III peptides from flavivirus envelope protein are useful antigens for serologic diagnosis and targets for immunization. Biologicals, 2010. 38(6): p. 613–8.

38. Babu, J.P., et al., Immunogenicity of a recombinant envelope domain III protein of dengue virus type-4 with various adjuvants in mice. Vaccine, 2008. 26(36): p. 4655–63.

## References

1. Arnaudov, L.N. and R. de Vries, Thermally Induced Fibrillar Aggregation of Hen Egg White Lysozyme. Biophysical Journal, 2005. 88(1): p. 515–526.

2. Kabir, M.G., M.M. Islam, and Y. Kuroda, Reversible association of proteins into sub-visible amorphous aggregates using short solubility controlling peptide tags. Biochim Biophys Acta Proteins Proteom, 2018. 2: p. 366–372.

